# EfNST: A composite scaling network of EfficientNet for improving spatial domain identification performance

**DOI:** 10.1101/2023.12.03.569798

**Authors:** Yanan Zhao, Chunshen Long, Na Yin, Zhihao Si, Wenjing Shang, Zhenxing Feng, Yongchun Zuo

## Abstract

Spatial Transcriptomics (ST) leverages Gene Expression Profiling while preserving Spatial Location and Histological Images, enabling it to provide new insights into tissue structure, tumor microenvironment, and biological development. The identification of spatial domains serves as not only the foundation for ST research but also a crucial step in various downstream analyses. However, accurately identifying spatial domains using computational methods remains a tremendous challenge due to the poor computational performance of many existing algorithms. Here, we propose EfNST, a deep learning algorithm based on a composite scaling network of the EfficientNet Network, designed specifically for the analysis of 10X Visium spatial transcriptomics data. We applied EfNST to three different datasets: human Dorsolateral Prefrontal Cortex, human breast cancer and mouse brain anterior. EfNST outperforms five advanced competing algorithms, achieving the best Adjusted Rand Index (ARI) scores of 0.554, 0.607, and 0.466, respectively. Notably, EfNST demonstrated high accuracy in identifying fine tissue structure and discovering corresponding marker genes with an improved running speed. In conclusion, EfNST offers a novel approach for inferring spatial organization of cells from discrete datapoints, facilitating the exploration of new insights in this field.

## INTRODUCTION

The single-cell transcriptomic^1^ has refined the research accuracy from the organizational level to the single cell, allowing researchers to analyze processes such as embryonic development, cell reprogramming, and disease occurrence with unprecedented resolution^2^. However, cell subpopulations act in an intertwined manner based on their specific spatial positions within a given tissue^3^. The spatial heterogeneity is a crucial characteristic for understanding organ function, cell fate regulation mechanism and cell lineage generation^4^. The single-cell transcriptomics technologies inevitably lose spatial information in the process of dissociating solid tissues into individual cells. Therefore, in order to better understand the spatial heterogeneity of different cells, it is necessary to simultaneously understand their transcriptional heterogeneity and spatial location information.

In recent years, the ST technologies has become a powerful tool in studying molecular biological mechanisms, which can obtain the spatial information along with transcriptomic expression profiling information of cells to analyze and characterize the expression profiles of specific cell types on a spatial scale. Therefore, the ST technologies has promising applications in oncology, immunology, developmental biology, neuroscience and pathology^5^. However, the low sensitivity, high dimensionality, multiple sparsity, high noise and multimodality of ST data pose significant challenges for dissecting cellular functions within the spatial context. Thus, it is a necessary to develop specialized computing tools based on ST data to accurately and robustly identify spatial domains and tissue structure.

Recently, many algorithms have been developed for identifying spatial domains by considering the similarities between neighboring spots to better explain spatial dependencies between the gene expression. Some of the algorithms are analogous to single cell processing approaches where spatial information is underutilized, such as BayesSpaces^6^, Giotto^7^, Seurat^8^, Scanpy^9^, stLearn^10^. Some other algorithms construct the deep autoencoder networks to learn low-dimensional embeddings of gene expression profiles and spatial information, and use embedding clustering to partition structural domains, these algorithms have their own limitations when it comes to leveraging transcriptomics, histological information, and spatial information, such as SpaGCN^11^, STAGATE^12^, SEDR^13^, CCST^14^.

In this study, we introduced the powerful convolutional neural network architecture of EfficientNet^15^ developed by Google to identify organizational spatial domains. It is an efficient, accurate, scalable, and adaptive to migration learning neural network architecture for a variety of visual tasks, and has better performance and efficiency on resource-limited devices. The algorithm incorporated deep learning network models while applying the EffectientNet network based on the combination of gene expression profiling data, spatial location and image data. On the human breast cancer dataset, EfNST accurately identifies subregions of tumors and can finely delineate tumor regions in comparison with existing methods. Similarly, on the other two datasets of human Dorsolateral Prefrontal Cortex and mouse brain anterior, it also performed a better recognition ability of organization spatial domains. The EfNST algorithm not only learned latent representations during implementation, but also showed a strong performance in downstream analysis.

## RRESULTS

### The EfNST framework

EfNST accurately identifies spatial domains by integrating Gene Expression Profiling, Spatial Location, and potential characterization of Histological Image information to elucidate heterogeneity in tissue structure (Fig. 1a-1d). Firstly, Fig. 1a performs preprocessing (see method) on the Gene Expression Profiling, Spatial Location and Hematoxylin and Eosin (H&E) image information in the ST datasets.

**Fig. 1.**
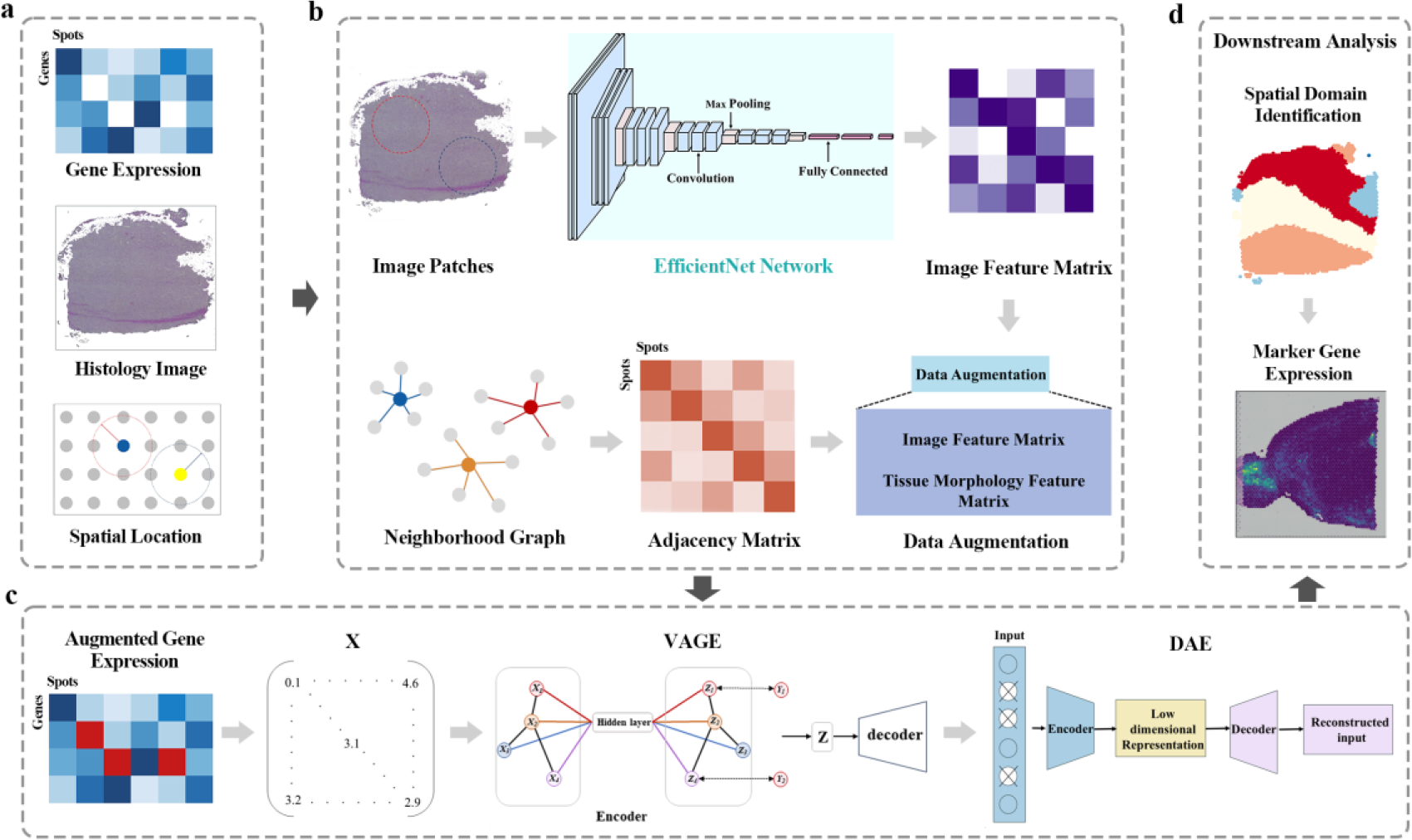
Workflow of the EfNST. (a) The input ST data are Gene Expression, Histological Images and Spatial Location; (b) EfNST processed the H&E Images and Spatial Locations to obtain Image Patches, which were processed using a pre-trained EfficientNet network to obtain Image Feature Matrix. Data Augmentation is performed for each spot based on the similarity of spots in spatial combined with the gene expression weights and the spatial location weights; (c) Final latent embedding is achieved using VAGE and DAE; (d) Latent representations can be used to perform Downstream Analysis.

Second, Image Features Matrix of the image is obtained by processing the histology image dataset using the EfficientNet network. The Tissue Morphology Feature Matrix is obtained by integrating the KNN algorithm to compute the Adjacency Matrix of neighboring spots. Augmentation the Feature Matrix through Data Augmentation (see method) (Fig. 1b). Then, the augmented gene expression matrix X is processed using the VAGE and the DAE, which can avoid the overfitting phenomenon caused by excessive features (Fig. 1c). Finally, the Low dimensional latent representations can be used to perform Downstream Analyses, such as identification of spatial domains and marker genes, and visualization (Fig. 1d).

### Application to the Dorsolateral Prefrontal Cortex dataset

To evaluate the basic performance of the EfNST algorithm, the DLPFC^16^ dataset consisting of 12 slices from the spatialLIBD database was analyzed. In the manually annotated region of the DLPFC, there are eight DLPFC slices with seven layers of organization. These slices span six cortical layers and white matter (WM), and the order of these layers is layer1 to layer6 to WM layer. The other four DLPFC slices have five layers of tissue structure, spanning four cortical layers and white matter layer in the order of layer 3, layer 6 and WM layer. The DLPFC dataset is chosen because the human Dorsolateral Prefrontal Cortex has clear and established morphologic boundaries and can be used as a benchmark dataset for comparison purposes.

Here, slides 151670 and 151507 were used to test the clustering performance of the EfNST algorithm. Slice 151670 in the DLPFC dataset is one of the few slices with five layers of organization (Fig. 2a). The manually annotated layers of slice 151670 was used for comparison, using ARI score to compare EfNST with the other five algorithms. EfNST achieved the highest ARI score of 0.554, which was 0.2 higher than the ARI value of SEDR. Compared to other methods, EfNST displays distinct layering and clearer boundaries, aligning more closely with manually annotated layer structures. However, when examining the spatial domains predicted by the Seurat and Scanpy algorithms, it was observed that most of the spots presented a mixed state with no discernible hierarchical structure (Fig. 2b).

**Fig. 2.**
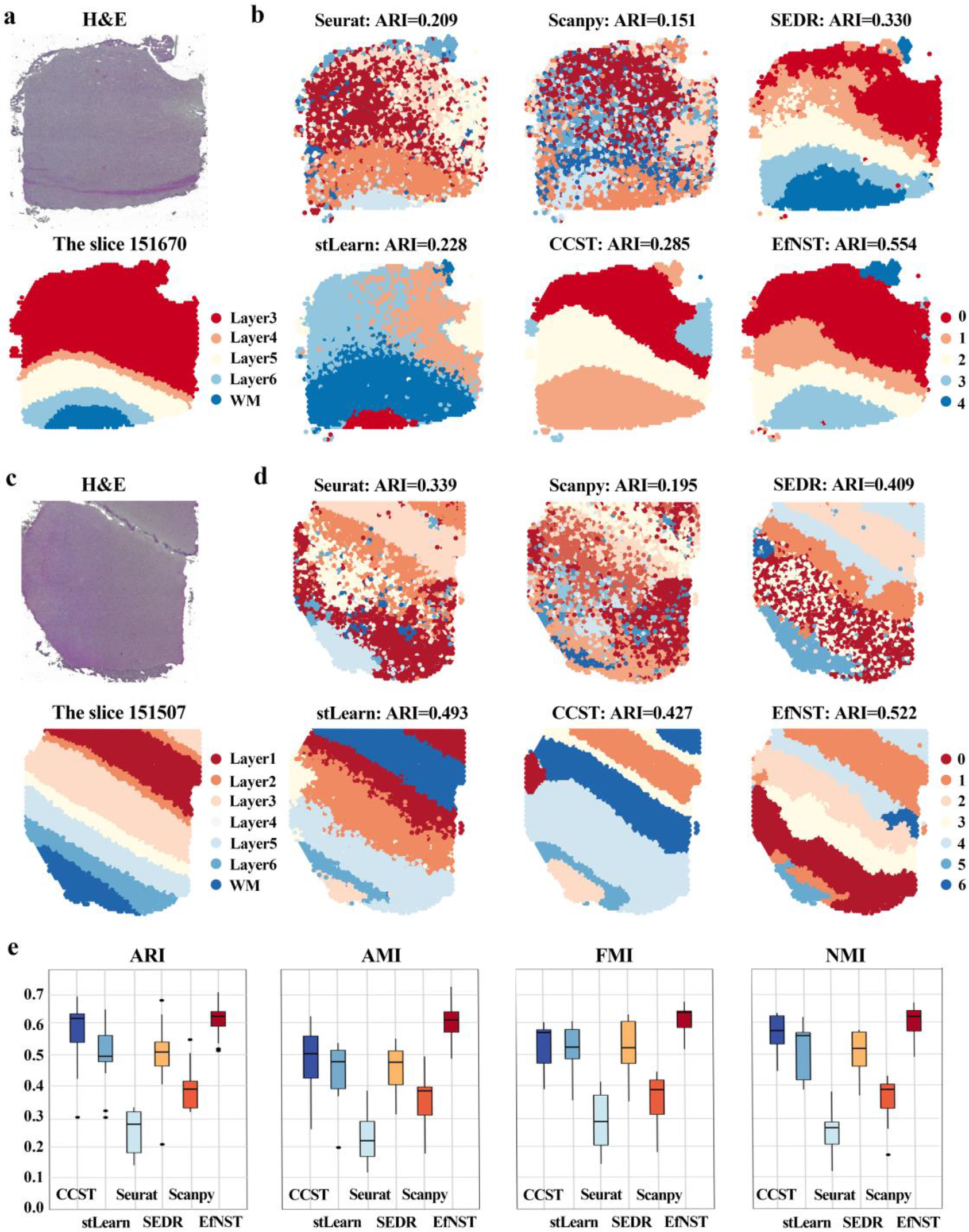
EfNST improves the ability to recognize the DLPFC layer structures. (a) H&E Images of slice 151670 and manually annotated layers; (b) Comparison of the performance of EfNST and existing state-of-the-art algorithms (Seurat, stLearn, Scanpy, CCST, SEDR) for recognizing laminar structures on slice 151670; (C) H&E Images of slice 151507 and manually annotated layers; (d) Comparison of the performance of EfNST and existing state-of-the-art algorithms (Seurat, stLearn, Scanpy, CCST, SEDR) for recognizing layer structures on slice 151507; (e) Boxplot comparing the performance of EfNST and existing state-of-the-art algorithms on 12 slices of the DLPFC. x-axis represents the six algorithms and y-axis represents the ARI, AMI, FMI, NMI scores, respectively.

Slice 151507 is the largest slice in the DLPFC dataset and has a seven-layer hierarchical structure (Fig. 2c). The manually annotated layers of the slice 151507 was used as the ground truth, and EfNST was compared with advanced algorithms, such as Seurat, CCST, Scanpy, SEDR and stLearn. We can observe that EfNST had the highest ARI score of 0.522, and the recognized spatial domains were accurate and clear (Fig. 2d).

To make a comprehensive evaluation of EfNST’s ability to recognize spatially structured domains, we used four clustered internal evaluation metrics to measure the similarity between all domains in all 12 slices and ground truth. Boxplot can be observed that EfNST achieved higher ARI, NMI, FMI and AMI scores in all slices in comparison with the shown 5 previous algorithms, demonstrating the better clustering performance and higher accuracy in identifying spatial domains of our EfNST algorithm (Fig. 2e).

### Application to the human breast cancer dataset

To tested the generalization ability of EfNST algorithm on tumor tissues, we also used the human breast cancer dataset with high differences between intra-tumor and inter-tumor. Strongly associated with poorer survival prospects, intra tumor heterogeneity in cancer is one of the main reasons of complicate treatment options. With a complex micro environment, tumor tissues exhibit highly heterogeneous^17^. Based on H&E image and manual pathologic labeling, the manually annotated regions were meticulously categorized into 20 regions to elucidate the difference of various regions on a subdivision scale in breast cancer tissue^18^. The 20 regions were further classified into four main types: ductal carcinoma in situ/lobular carcinoma in situ (DCIS/LCIS), healthy tissue, invasive ductal carcinoma and tumor edge (Fig. 3a).

**Fig. 3.**
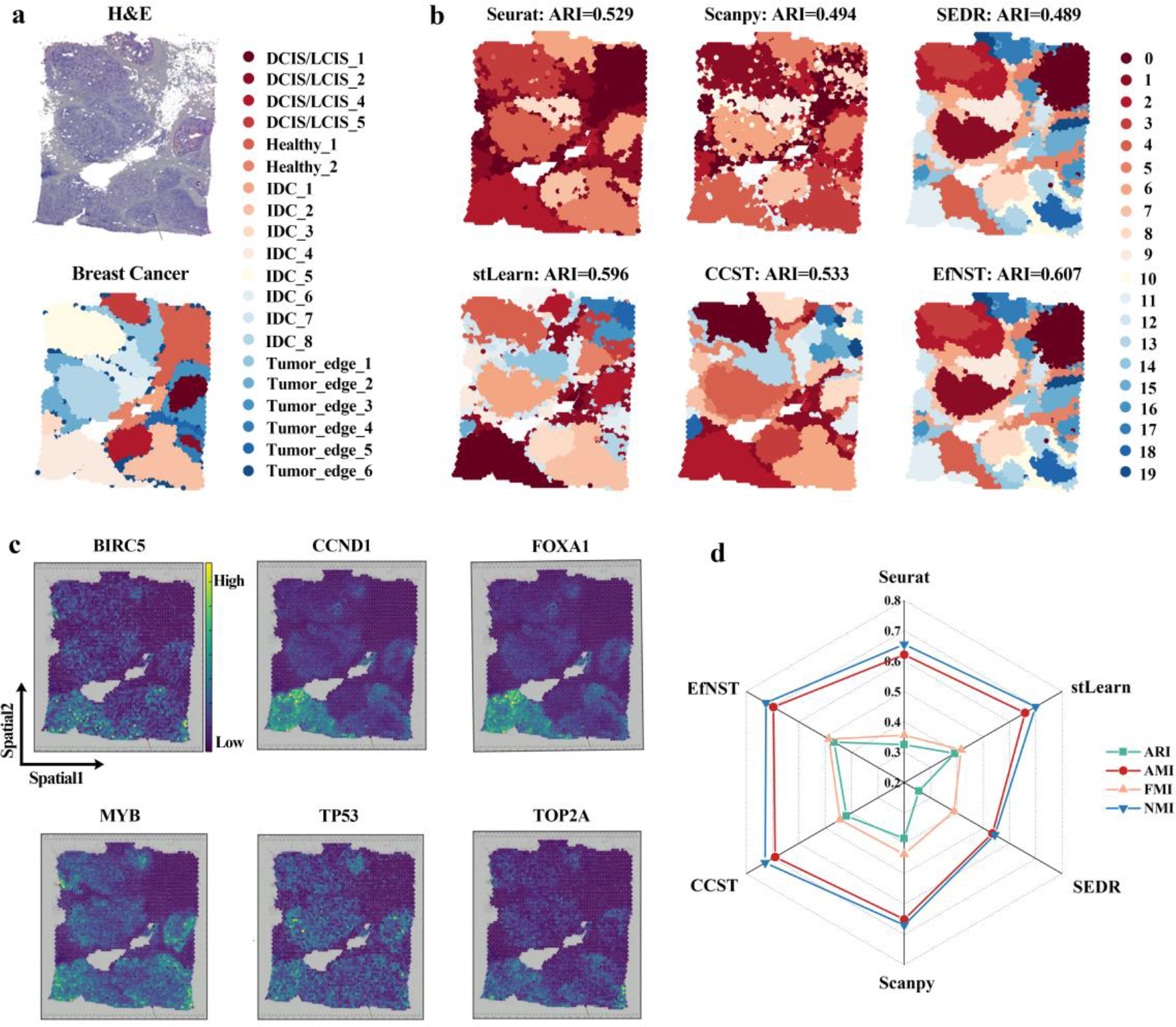
EfNST can recognize the spatial domain of human breast cancer with a finer level of granularity. (a) H&E Images of human breast cancer sample (Block A, Section 1) and manually annotated regions; (b) Performance comparison of using EfNST and existing state-of-the-art algorithms (Seurat, stLearn, Scanpy, CCST, SEDR) to recognize spatial domains of breast cancer tissue; (c) Spatial expression distribution of breast cancer marker genes; (d) Radar chart comparing the clustering performance of EfNST with existing state-of-the-art algorithms in the breast cancer. Axes represent scores.

The clustering comparisons of EfNST with the other five algorithms on the breast cancer dataset were illustrated in a clustering plot. While at the clustering level, many of the predicted clusters in Seurat and Scanpy were fragmented and discontinuous. On the contrary, the CCST, stLearn, SEDR and EfNST algorithms were able to generate more independent clusters with a better clustering performance. The same result was observed in the ARI score, where EfNST had the highest ARI score of 0.607, while SEDR, Scanpy had lower scores around 0.48. CCST, stLearn and SEDR received higher ratings, ranging between 0.52-0.59(Fig. 3b). Compared to the other methods, EfNST demonstrated higher scores in all four clustering internal metrics (Fig. 3d). In order to test the ability of EfNST to detect the biological correlation of tissue structure, we further observed the spatial expression distribution of known breast cancer marker genes (e.g., FOXA1^19^, TP53^20^, BIRC5^21^, TOP2A^22^, CCND1^23^, MYB^24^). The results showed all of the marker genes can be clearly identified (Fig. 3c).

### Delineating spatial domains on the complex mouse brain anterior dataset

In neuroscience research, the tissue spatial structure of mouse brain garnered significant attention. The strong similarity of the structure and function of brain tissue between the mouse and human make it an ideal model for studying brain diseases and cognitive functions. The EfNST algorithm was employed for cluster analysis of mouse brain tissue based on H&E images. A total of 52 regions in the anterior part of the mouse brain were manually annotated and segmented^25^ (Fig. 4a).

**Fig. 4.**
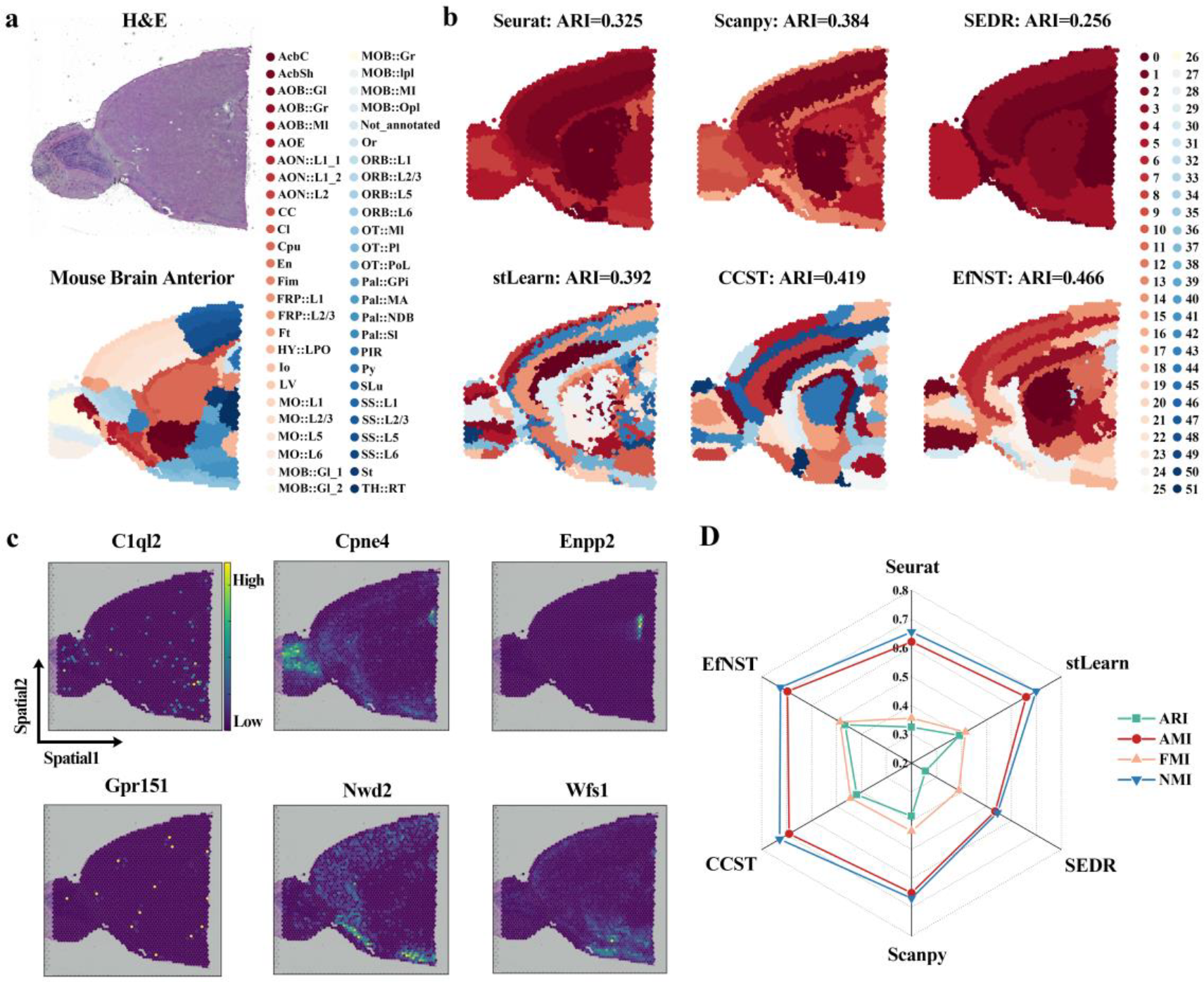
EfNST can recognize complex spatial domains in the Mouse Brain Anterior. (a) H&E Images of mouse brain anterior and manually annotated regions; (b) Performance comparison of using EfNST and existing state-of-the-art algorithms (Seurat, stLearn, Scanpy, CCST, SEDR) to recognize spatial domains of mouse brain anterior tissue; (c) Spatial expression distribution of mouse brain anterior marker genes; (d) Radar chart comparing the clustering performance of EfNST with existing state-of-the-art algorithms in the mouse brain anterior. Axes represent scores.

The ability of EfNST to recognize spatial domains assessed against five other algorithms using data of the mouse brain anterior. The EfNST demonstrates the ability to identify concentrated areas with clear boundaries. The clustering performance of six algorithms (i.e., EfNST, Seurat, CCST, Scanpy, SEDR and stLearn) were evaluated using four internal clustering evaluation metrics. It was found that EfNST achieved the highest ARI score of 0.466, while the ARI scores of other algorithms were below 0.4 (Fig. 4b). In addition, the similar trend was also observed in the other three internal indicators (Fig. 4d). To further validate the biological relevance of EfNST in detecting these structures, marker genes were detected in the anterior part of the mouse brain. In Fig. 4c, the genes Enpp2^26^, Nwd2^27^, Cpne4^28^ and C1ql2^29^ were found to be highly expressed in this tissue section. It has been proved that Wfs1^30^ is a potential target for regulating tau protein in the treatment of Alzheimer’s disease and other neurodegenerative diseases. Gpr151^31^, a highly expressed gene, has been experimentally shown to play a key role in the pathogenesis of neuropathic pain in injured DRG neurons and may be a potential target for the treatment of neuropathic pain. Additionally, EfNST also possesses a significant advantage in terms of computational speed (Fig. 5). These results showed that EfNST can accurately identified spatial domains and verified their biological correlations.

**Fig. 5.**
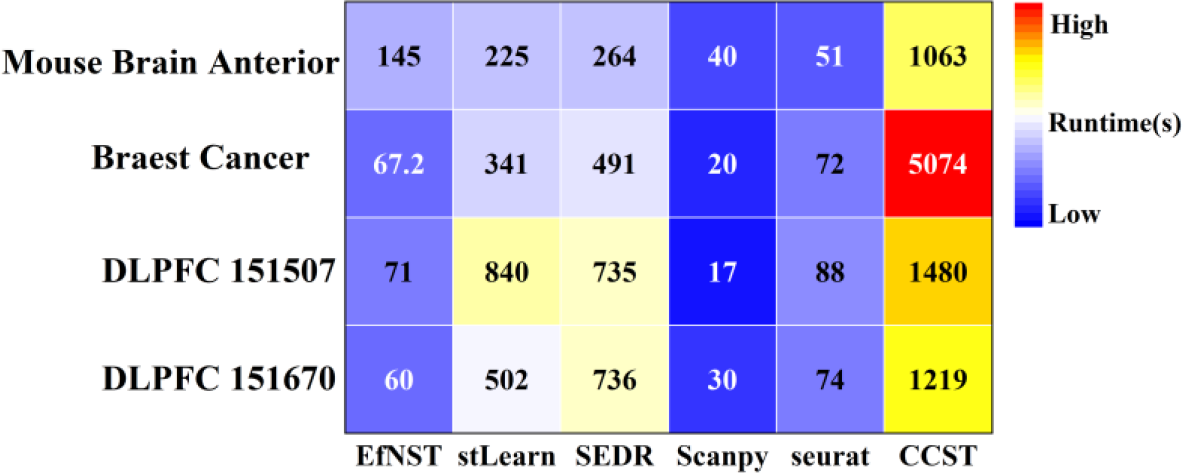
Compared to other algorithms in the article, the running speed(s) of EfNST has been improved.

## DISCUSSION

Currently, identifying the spatial domains of tissue structures is a critical step in resolving the regulation mechanism of key genes in tumor biology, developmental biology and neurobiology. In this study, we introduced the EfNST algorithm, a precise and efficient approach for accurately identifying spatial domains by integrating gene expression profiles, spatial localization, and histological images. Compared to previous multi-algorithms, EfNST not only improved the ability to identify the spatial domains of tissues, but also dissected the spatial domains of cancerous and normal tissues at a fine scale on three 10x Visium datasets of DLPFC, breast cancer and mouse brain anterior. The latent representations learned by EfNST can be effectively used in downstream analyses including clustering, visualization and marker genes detection. Additionally, by incorporating efficient network architectures such as EfficientNet, VAGE, and DAE, we can enhance the speed of execution and data augmentation. EfNST provides a robust and precise computational framework for researchers to analyze spatial domains within tissues.

## METHODS

### Datasets and preprocessing

To evaluate the performance of EfNST, we downloaded three types of 10x genomics ST datasets (https://www.10x-genomics.com)which were commonly used in previous works. (1) The benchmark dataset of human Dorsolateral Prefrontal Cortex consists of 12 slices. (2) Human breast cancer dataset (3798 spots in tissue section and the number of median gene is 6026 in each site). (3) Mouse brain anterior dataset (2696 spots in tissue section and the number of median gene is 6015 for each site). In the data preprocessing phase, we performed Quality Control, Log transformation and Normalization of gene expression data, the Principal Component Analysis (PCA) was used to reduce the dimension of gene expression profiling data in the ST datasets.

### EfficientNet

Tan and Le proposed the EfficientNet^15^ of the convolutional network from the idea of augmenting the network in a principled way to obtain higher accuracy in 2019.The difference between EfficientNet and the previously designed networks were that it adopted a unified method of expanding the depth, width, and resolution of the network. EfficientNet is developed as a new baseline network for ensure the effectiveness of the network. On the basis of not changing the predefined operators in the baseline network, it scales the network length, width, and resolution in a fixed proportion to achieve maximum model accuracy. The corresponding mathematical formula are as follows:

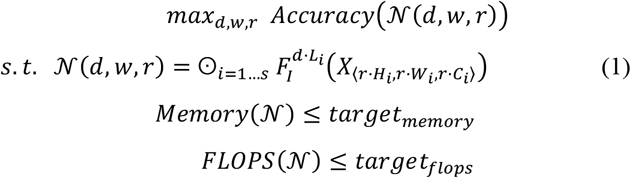

Where d,w and r are coefficients used to scale the depth, width and resolution of the network. *F*_*i*_, *L*_*i*_, ⟨*H*_*i*_ *,W*_*i*_⟩and *C*_*i*_ represent Operator, Layers, the resolution of the input and Channels, where i represents the Stage.

Tan and Le founded that the optimized *d, w* and *r* were interdependent, so they proposed a new uniform scaling method for *d, w* and *r*, which was a necessary requirement for achieving better accuracy and efficiency. The method selects the composite coefficient *ϕ* scale the network width, depth and resolution uniformly in a principled manner. The corresponding mathematical formula are as follows:

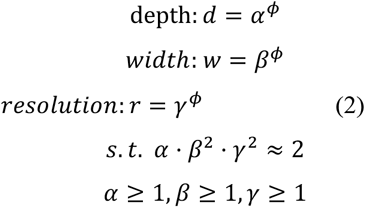

where d,w and r are dynamic coefficients used to control depth, width and resolution, and *ϕ* is a constant used to control model complexity.

The series of models obtained after expanding the EfficientNet baseline network were called EfficientNets. Compared to previous ConvNets, EfficientNets have higher accuracy and efficiency. In comparison with other popular deep learning frameworks, such as ResNet^32^, Inception^33^ and MobileNet^34^, EfficientNets can achieve better computational performance. In addition, EfficientNets also show the best performance in image processing compared to other image classification benchmarks, such as ImageNet^35^ and CIFAR-100^36^.

At the same time, EfficientNets can better adapt to robust to data expansion and noise while improving accuracy and efficiency. Thus, it is more suitable for practical applications. Currently, the EfficientNes are popularly used in biomedical aspects, such as breast cancer detection^37^, retinal disease screening^38^ and brain tumor classification^39^.

### Data Augmentation

Data augmentation^40^ is widely used in deep learning to expand the training dataset, increasing data diversity and improving the algorithm’s generalization and robustness. This, in turn, enhances performance. Cropped image data is combined with the original data to expand gene expression data, thereby increasing the number of training datasets.

### Denoising Autoencoder

The DAE^41^ can avoid losing useful information after inputting the original data by reconstructing the input data containing noise, and learn better feature representations. The features it learned are almost identical to those learned from data without superimposed noise and more robust.

### Variational Graph Autoencoder

The VAGE^42^ was proposed by Kipf and Welling utilized latent variables to learn latent representations in graph structured data based on VAE. According to the given unweighted undirected graph G and its adjacency matrix A, it uses the latent variable Zi to reconstruct the Adjacency matrix:

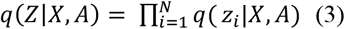

X is the feature matrix of the nodes; A is the adjacency matrix:

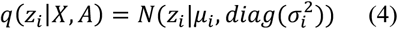

where *µ* = *GCNµ*(*X, A*) is the mean of the feature vector; *logσ* = *GCNσ*(*X, A*) is the variance of the node vector.

The defined two-layer neural network is:

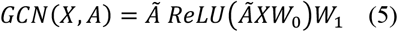

where *GCN*_*µ*_(*X, A*) and *GCN*_*σ*_(*X, A*)share the first level parameter W_0and do not share the second level parameter *W*_1_;

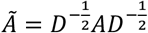 is the symmetric normalized adjacency matrix.

After this, the decoder reconstructs the adjacency matrix using the inner product of the latent variables.

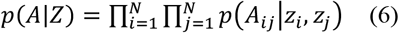

Among them 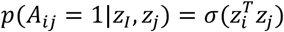,

The KL scatter is added to the loss function to improve the generalization ability and robustness of the model:

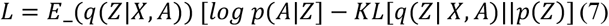

Where *p*(*Z*) = Π_*i*_ *N*(*z*_*i*_|0, *I*).

### Clustering

Based on the VAGE embedding, EfNST identifies a set of spots with similar transcriptomes clustered in space by a clustering algorithm. We used the Leiden method in the Scanpy package for clustering, which was more reasonable in clustering, results and may result in more subgroups, and runed faster.

### Evaluation indicators

We utilized the Scikit-learn^43^ package for calculating the clustering evaluation metrics. We had ground truth files of all three datasets and we employed four clustering evaluation metrics: Adjusted Rand Index (ARI), Adjusted Mutual Info (AMI), Fowlkes-Mallows Index (FMI) and Normalized Mutual Info (NMI)^44^. The metrics are used to evaluate the algorit-hm’s ability to identify spatial domains, ARI is a main metric for measuring clustering ability.

The formula for the evaluation metrics is as follows:

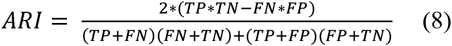

Where TP, TN, FP and FN represent true positive, true negative, false positive and false negative. The value range of ARI is [0,1], a higher ARI score indicates a stronger consiste-ncy between the clustering results and the ground truth.

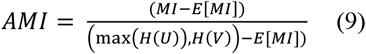

MI denotes mutual information, H(U) and H(V) represent the entropy of the clustering result and the ground truth, respe-ctively. Meanwhile, E[MI] denotes the expected mutual infor-mation with random assignment. A higher AMI score indicat-es a stronger clustering ability of the algorithm.

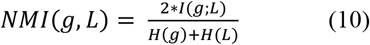

The variable *g, L*, and *I*(*g; L*) represent the ground truth, the result of clustering, the Cross Entropy and the mutual information. The mutual information is calculated using the formula *I*(*g; L*) = *H*(*g*) − *H*(*g*|*L*).

Fowlkes-Mallows Index (FMI) is commonly used to measure the similarity between clustering and the ground truth in clustering, its score range is [0,1].

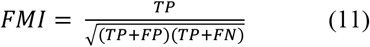

Where *TP, FP* and *FP* respectively represent the number of true positives, the number of false positives and false negative. A higher FMI score indicates a good similarity between the clusters and the ground truth.

## Data availability

The three test datasets used in this study are available at https://www.10x-genomics.com. These datasets include: human Dorsolateral Prefrontal Cortex, Human breast cancer dataset and Mouse brain anterior dataset.

